# The tiger shark, *Galeocerdo cuvier* (Péron & Lesueur, 1822), genome reveals genomic and epigenomic signatures of shark evolution and demographic history

**DOI:** 10.64898/2026.11.05.764724

**Authors:** Jessica R. Pearce, Adrianne Doran, Emma De Jong, Sarah Kwong, Oliver Jewell, Ben D’Antonio, Matthew W. Fraser, Shannon Corrigan, Luke Thomas, Rich Edwards

## Abstract

Cartilaginous fishes (Chondrichthyes) represent one of the oldest extant vertebrate lineages, persisting for over 450 million years, yet genomic resources remain limited for many species despite widespread conservation concern. The tiger shark, *Galeocerdo cuvier* (Péron & Lesueur, 1822), the sole surviving member of the family Galeocerdonidae, is a circumglobally cosmopolitan predator vital in ecosystem regulation but is experiencing population declines. Here, we generated the first chromosome-scale haplotype-phased reference genome for the tiger shark and integrated comparative and population genomics with methylomics to characterise its genome architecture and evolutionary history. This assembly presented the first evidence of male heterogamety in this species. Comparative analyses revealed lineage-specific transposable element expansions across the chondrichthyan class suggesting ongoing genome restructuring in Carcharhiniform sharks. We report the first hydroxymethylome in a chondrichthyan, higher than that of other vertebrates, hinting at novel DNA evolutionary and regulatory pathways in an early-diverging lineage of jawed vertebrates. We identified genomic signatures of prolonged population decline that correlated with historical climate fluctuations and have resulted in sustained low levels of genetic heterozygosity in this species. These findings offer new insights into shark and broader vertebrate genomic and epigenomic evolution while providing valuable resources for conservation genomics of one of the world’s most widespread top-order predators in the face of accelerating environmental change.

## 1. INTRODUCTION

Cartilaginous fish of the class Chondrichthyes (sharks, rays, skates and chimaeras) are an early-diverging (> 450 million years ago) lineage of jawed vertebrates. Chondrichthyan evolutionary history has been shaped by repeated episodes of diversification and extinction, with substantial lineage turnover underlying their remarkable persistence (Kocáková et al., 2025). This history is reflected in a contemporary phylogeny that is characterised both by sparsely populated longer branches and species-rich radiations. Today, these surviving lineages face widespread anthropogenic pressures: recent global assessments estimate that about a third of chondrichthyan species are threatened with extinction, primarily due to overexploitation (Dulvy et al., 2021; Kim et al., 2025; Schnetz et al., 2024). Deep evolutionary history and contemporary vulnerability together underscore their conservation importance as a group harbouring both phylogenetic and ecological diversity. Tiger sharks, *Galeocerdo cuvier* (Péron & Lesueur, 1822), are a large-bodied (up to ∼ 5 m) cosmopolitan species, distributed circumglobally from the tropics to warm-temperate marine waters (Ferreira et al., 2015; Heithaus et al., 2007). They play vital roles in maintaining ecosystem health and function through top-down regulation (Hammerschlag et al., 2015; Heithaus et al., 2012) and are a potential keystone species in some areas (Ferreira et al., 2017; Heithaus, 2001). The tiger shark lineage has experienced repeated global disturbance events and geologically rapid climate shifts over the course of their ∼ 13.8 million year trajectory (Türtscher et al., 2021), and *G. cuvier* is the last extant member of the family Galeocerdonidae. In the Anthropocene, this species distribution also suggests sensitivities to ocean conditions like salinity (D’Antonio et al., 2025) and temperature (Hammerschlag et al., 2022), demonstrating a potential vulnerability to contemporary global change that risks loss of phylogenetic uniqueness. In 2019, tiger sharks were assessed as Near Threatened according to the International Union for the Conservation Nature Red List criteria due to global population declines approaching 30% and anticipated further reductions in light of ongoing exploitation practices (Ferreira & Simpfendorfer, 2019; Manuzzi et al., 2022).

A single high-quality representative genome assembly (i.e. reference genome) presents a powerful scientific resource that can be used to derive insights about species biology, ecology and evolutionary history, including information that may be relevant to conservation and management (De León et al., 2023; Ellegren, 2014), such as ancient effective population sizes (N_e_) and demographic change over time (Hohenlohe et al., 2021; Matz, 2018; Shafer et al., 2015), or runs of homozygosity (ROH) across the genome that indicate the degree and timescale of inbreeding (Robinson et al., 2019; Saremi et al., 2019; Stanhope et al., 2023). These metrics can provide insight into species resilience, where historical demographic changes can be linked to past environmental events, and potential responses to future threats inferred.

Methylation at CpG dinucleotides is a widespread epigenetic feature of vertebrate genomes, playing a key role in regulating gene expression, maintaining genome stability, and suppressing transposable element activity (Bird, 2002; Slotkin & Martienssen, 2007; Suzuki & Bird, 2008). While much of the vertebrate genome is highly methylated, CpG methylation level varies across genomic features, with regulatory regions such as promoters often exhibiting hypomethylation which is generally associated with active transcription (Varriale, 2014). Recent advances in long-read sequencing provide new opportunities to characterise these patterns at genome-wide scale, as native DNA sequencing can simultaneously resolve genomic sequence and directly detect DNA modifications. As an early-diverging vertebrate lineage, chondrichthyans provide an important comparative system for understanding the evolution of vertebrate epigenetic regulation (Varriale, 2014). Yet only a handful of other studies have investigated methylation levels and patterns within the Chondrichthyan class (Beal et al., 2022; Beal et al., 2021; Bock et al., 2026; Colwell et al., 2018; de Jong et al., 2026; Peat et al., 2017).

Given its distinctive phylogenetic and ecological significance, the lack of genomic resources for the tiger shark represents a limitation to research required to inform its conservation and management. Combining multiple sequencing platforms, we generated and characterised the first chromosome-level, haplotype phased reference genome for the tiger shark, providing a comprehensive view of its genome architecture, sex chromosomes and methylation landscape. We use this genomic resource to investigate the evolutionary and demographic history of this enigmatic species, providing insights into its past and potential future resilience. We also report the first 5-hydroxymethylcytosine (5hmC) methylome in a chondrichthyan species. Together, these findings provide a valuable genomic resource for understanding the evolution, biology, and conservation of this ecologically important predator.

## 2. METHODS

### 2.1 Genome sequencing

#### 2.1.1 Sample collection

To create the tiger shark reference genome, whole blood and tissue samples were obtained from a male individual caught in Exmouth Gulf, Western Australia, in 2023 as per procedures approved by The University of Western Australia’s Animal Ethics Committee (approval number 2021/ET000634), under a fishing exception (number 251157423) granted by the Department of Primary Industries and Regional Development (DPIRD). Whole blood was immediately transferred into 100% molecular grade ethanol in a 1:10 ratio and flash frozen in liquid nitrogen onboard the research vessel. To assess for heteromorphic sex chromosomes, another four tiger sharks (two males and two females) were sampled from this location in 2025, where tissue biopsies were taken from pectoral fins and flash frozen in liquid nitrogen. All samples were stored at -80C upon return to laboratory facilities until processing.

#### 2.1.2 DNA extraction and preparation

The Nanobind CBB kit (PacBio, Menlo Park, CA, USA) and nucleated blood procedure was used to extract high molecular weight (HMW) DNA from the blood sample for PacBio HiFi (high-fidelity long-read) sequencing. A 100µl aliquot of the preserved blood sample was combined with 100µl of 1 X Phosphate Buffered Saline (PBS) for each extraction. Multiple HMW DNA extractions were performed from the same individual to obtain sufficient DNA yield required for sequencing. DNA concentration of the extracted genomic DNA (gDNA) was initially tested with the Qubit dsDNA BR (Broad Range) kit on a Qubit 4 Fluorometer (Thermo Fisher Scientific Inc., Waltham, MA, USA) before proceeding with a 1X PacBio SMRTbell bead clean-up of each sub-sample. The Qubit dsDNA HS (High Sensitivity) and Thermo Scientific™ NanoDrop™ One Microvolume UV-Vis Spectrophotometer (Thermo Fisher Scientific Inc.) were then used to determine final concentration of DNA as well as A260/A280 and A260/A230 purity ratios. Fragment length profiles were then assessed using a Femto Pulse System with a 165kb kit (Agilent Technologies, Santa Clara, CA, USA).

#### 2.1.3 PacBio DNA library preparation and sequencing

Post bead clean-up, the HMW DNA sub-sample with the largest fragments was divided in half and mechanically sheared with a Megarupter 3 (30 speed) to shear ultralong DNA fragments targeting 18 kbp. The gDNA was then cleaned using 1X PacBio SMRTbell beads, and fragment length and concentration were checked with the Femto Pulse and Qubit dsDNA HS kit again respectively. The PacBio short-read eliminator (SRE) XS kit was used to deplete short DNA fragments up to 10 kb. The quality control process of 1X PacBio SMRTbell bead clean-up, Qubit dsDNA HS and Femto Pulse was performed again. The final fragment length distribution of each library was centred around 19,942 and 17,320 kbp. Libraries were prepared using the PacBio SMRTbell prep kit 3.0. Final DNA library concentrations were checked with the Qubit dsDNA HS kit and Femto Pulse before proceeding with sequencing preparation. Both sub-samples were prepared for HiFi sequencing with the PacBio Revio polymerase kit (two sequencing libraries), and sequenced across three SMRT Cells on the PacBio Revio platform.

#### 2.1.4 Hi-C DNA library preparation and sequencing

Short-read Hi-C sequencing was used to detect genome-wide chromatin interactions and provide long-range linkage information to scaffold, order, and orient contigs generated from long-read data (Matz, 2018). The Dovetail Omni-C kit and Non-mammalian Samples Protocol version 1.2B for nucleated blood was utilised for Hi-C lysate extraction with 200µL of 1:10 blood in ethanol. The resulting digestion profile was assessed using the High Sensitivity D5000 ScreenTape assay on a 4200 TapeStation System (Agilent Technologies) and concentration checked using the Qubit dsDNA BR kit on a Qubit 4 fluorometer before proceeding with the proximity ligation using Dovetail Proximity Ligation Core. The DNA Hi-C library was prepared with the Dovetail® Library Module for Illumina® kit. Ligation capture was performed with the Dovetail® Dual Index Primer Set #1 for Illumina and the final library quantified with a Qubit dsDNA HS kit on a Qubit 4 fluorometer and fragment size distribution was assessed again using a TapeStation. The library was initially sequenced on an Illumina (San Diego, CA, United States) iSeq100 to assess library complexity against an assembly of the HiFi data, before being sequenced to 60x coverage on an Ilumina NovaSeq 6000 System using an S1 flow cell generating 2 x 150 bp paired-end reads.

#### 2.1.5 Nanopore DNA library preparation and sequencing

Following the HMW DNA extraction and bead clean-up, library preparation was conducted with the SQK-ULK114 (Ultra-Long DNA) sequencing kit (Oxford Nanopore Technologies (ONT), Oxford, United Kingdom). This was sequenced on one Flow Cell over 48 hours on the ONT PromethION24 device. A DNase wash and library reload was performed additionally at 24 hours to help unblock pores and increase overall output.

#### 2.1.6 Illumina draft genome DNA library preparation and sequencing

The four fin tissue biopsy samples were extracted using the DNeasy Blood and Tissue kit (Qiagen) following the standard protocol with a 3-hour tissue digest. gDNA concentration was measured using the Qubit dsDNA BR kit. Following the DNA tissue extraction, DNA libraries were prepared using Illumina DNA Prep kit with 150 ng input following manufacture’s guidelines. These samples were then sequenced at 50X depth per sample on a NovaSeq 6000 System using an S2 flow cell generating 2 x 150 bp paired-end reads.

### 2.2 Genome assembly, curation and annotation

A chromosome-level reference genome was assembled following the Ocean Genomes publicly available pipeline (Parata, Anstiss, et al., 2025; Parata, de Jong, et al., 2025) (https://github.com/Minderoo-OceanOmics-Centre-UWA/OceanOmics-OceanGenomes-ref-genomes), which is based on the Vertebrate Genomes Project workflows (Larivière et al., 2024; Rhie et al., 2021). Package version control was implemented using Singularity (v3.11.4) containers. Initial quality control of raw HiFi reads to filter out adapter contamination was done with HiFiAdapterFilt (v2.0) (Sim et al., 2022). Meryl (Rhie et al., 2020) was used to construct a k-mer database and histogram for estimating genome size with GenomeScope2 (Ranallo-Benavidez et al., 2020).

A preliminary genome assembly was created de novo with HiFiasm v0.19.8 (Cheng et al., 2021) to assess Hi-C library complexity before Hi-C data was integrated. After the Hi-C library was determined to pass quality control measures, HiFiasm was run again to assemble contigs while integrating the Hi-C and Ultra-long ONT data to create the phased contig assembly and assembly graphs. Genome assembly statistics were summarised with gfastats v1.3.6 (Formenti et al., 2022). Genome completeness, both k-mer based and gene-based, was assessed with Merqury v1.3 (Rhie et al., 2020) and Benchmarking Universal Single-copy Orthologs (BUSCO v5.6.1 vertebrata_odb10 database; (Manni et al., 2021) respectively. The Hi-C reads were mapped to assembly contigs, using bwa (Vasimuddin et al., 2019), pairtools and samtools v1.16.1 (Danecek et al., 2021), according to the Dovetail Genomics Omni-C documentation (https://omni-c.readthedocs.io/en/latest/index.html). Chromosome-scale scaffolds were constructed with YaHS (Zhou et al., 2023). Decontamination of the genome assembly was done with the NCBI Foreign Contamination Screen to remove other species genetic sequences, then Tiara v 1.0.3 (Karlicki et al., 2022) to identify any other contamination in the assembly, which were then removed with BBMap v 39.01 (Bushnell, 2014). Manual assembly curation was conducted with PretextView (Harry, 2023) to visualise the scaffolds. Hi-C contacts were utilised to inform chromosome assignment, scaffolding and correction of mis-assemblies. Prior to this, bedgraph coverage and gaps tracks were generated with minimap2 v2.26 (Li, 2018), samtools and bedtools v2.30.0 (Quinlan, 2014). Assembly statistics, quality and completeness were assessed for the final curated haplotypes using gfastats, Merqury and BUSCO.

RepeatModeler v2.0.5 was used to build a de novo repeat library, which was then utilised by RepeatMasker v4.1.5, run with NCBI/RMBLAST v2.14.1 to soft mask the genome (Green, 2015; Hubley, 2015; Smit, 2004) and assess repeat content. The soft masked assembly was then annotated with GeMoMa v1.9 (Keilwagen et al., 2019) using the RefSeq assembly and gff annotation for *Scyliorhinus canicula* (smaller spotted catshark; NCBI accession: GCF_902713615.2). Windows of 500Kbp were generated and used to calculate GC content, gene density and repeat density across all chromosomes. Results were visualised using the circlize R package v0.4.18 (Gu et al., 2014).

### 2.3 Sex chromosome identification

The raw fastq files form the four high-coverage draft genome samples were filtered with fastp (Chen et al., 2018) to eliminate poor quality reads. These were aligned to the reference haplotype 1 assembly with bwa-mem2 v2.2.1 (Vasimuddin et al., 2019). Samtools v1.16.1 (Danecek et al., 2021) was then used to filter, sort and index the resulting bam files (--min-MQ 10). Duplicate reads were removed with the MarkDuplicates function (--REMOVE_DUPLICATES true) from the Picard package within GATK4 v4.2.5.0 (McKenna et al., 2010). Samtools depth function (-a) was used to assess the sequence depth per base, which was then averaged per chromosome for each sample (Yamaguchi et al., 2023). The mean depth per chromosome was calculated for the male and female samples separately and compared in R (Allaire, 2012).

### 2.4 Methylation analysis

Raw PacBio HiFi bam files were first aligned back to the curated tiger shark reference assembly using PacBio’s minimap2 SMRT wrapper for PacBio data, pbmm2, through Bioconda, with the preset CCS. Methylation probabilities were called from the alignment with PacBio’s pb-CpG-tools and aligned_bam_to_cpg_scores function. Only sites with a minimum of 10X coverage were retained from the resulting bed output. Site methylation level was calculated as the proportion of modified reads called. Genome-wide methylation was calculated as the sum of modified reads across all sites divided by the sum of valid reads across all sites. Average methylation across 500Kbp windows was calculated and visualised within the circos plot. Global methylation was additionally calculated separately for gene, promoter and intergenic regions based on genomic coordinates extracted from the genome annotation.

ONT basecalling was conducted with dorado v1.4.0 (Oxford Nanopore Technologies) to produce methylation tags for both 5-Methylcytosine (5mC) and 5-Hydroxymethylcytosine (5hmC) modified bases using the high accuracy model (hac) and a minimum average read qscore of 7. This model was run twice to call both CpG associated methylation (5mCG_5hmCG) and all-context (5mC_5hmC). Methylation calls were extracted from the bam file using ONT’s Modkit v0.6.1 with default pileup for all-context methylation. To assess CpG methylation, however, calls were restricted to CpG sites and base modification counts were summed across strands as CpG is palindromic. Alignment to the reference assembly was done within the dorado basecaller and again to realign CpG position within Modkit. Outputs were filtered to only retain either 5mC or 5hmC modified sites and impact of filtering for sequencing depth was tested at 5X and 10X thresholds. Global, windowed and feature level coverage was again calculated as per the HiFi methylation above. Potential associations between 5mCG methylation, gene content and repeat density were tested within each chromosome using Spearman’s rank correlations across 100 kb windows, with significance assessed using a circular-shift permutation test (5,000 permutations) to account for spatial autocorrelation between adjacent windows.

### 2.5 Comparative genomic analyses

Seven other Chondrichthyan species (*Callorhinchus milii*, *Pristis pectinata*, *Chiloscyllium punctatum*, *Carcharodon carcharias*, *Scyliorhinus canicula*, *Carcharhinus amblyrhynchos* and *Sphyrna mokarran*) with publicly available reference genome assemblies were selected to generate a phylogenetic tree and compare against the novel tiger shark assembly (Accession and citation details for all assemblies: Table S1). Of these, 6 species had available annotations on RefSeq. The great hammerhead shark (*Sphyrna mokarran*) did not have any annotation information available at the time of publication, and so was generated for this study following the same protocol as for the novel tiger shark assembly described previously. The soft masked assembly from RepeatMasker was annotated with GeMoMa using the RefSeq assembly and gff annotation for *Scyliorhinus canicula*. This was the most closely related species with an available annotation for both the tiger and great hammerhead sharks (same order: Carcharhiniformes). Repeat content was assessed for the genomes of all species as above using RepeatModler and RepeatMasker. Divergence landscapes were calculated from the output, classified by repeat class and plotted in R, with recently diverged repeats (<5%) highlighted. The significance of recent repeat class expansion was statistically tested in R through an overrepresentation test.

Phylogenies were reconstructed for these eight species from single-copy orthologues identified through OrthoFinder v2.5.5 (Emms & Kelly, 2019) with the annotation protein fasta files used as input. Sequence alignment was conducted within OrthoFinder with the -M msa option. The resulting single-copy orthologue alignments were extracted and trimmed with trimAL v1.5 (Capella-Gutiérrez et al., 2009) using -automated1 which is optimised for maximum likelihood phylogenetic tree reconstruction. These sequences were then concatenated into a supermatrix and partitions generated with AMAS (Borowiec, 2016). A maximum likelihood tree was built with IQtree2 v2.2.2.6 (Minh et al., 2020) using the partitions with 1000 Ultra-fast bootstrap replicates and the model ‘MFP+MERGE’.

### 2.6 Demographic history

A pairwise sequentially Markovian coalescent (PSMC) method (a coalescent-based hidden Markov model) was conducted to accurately infer ancient population history from a single high coverage genome (Ellegren, 2014; Sheehan et al., 2013). First, the filtered PacBio HiFi reads were realigned back to the reference assembly haplotypes 1 and 2 separately using Minimap2 v2.26 (Li, 2018) (-x map-hifi-a), which was then converted to a bam file and quality filtered (--min-MQ 10), sorted and indexed with Samtools v1.16.1 (Danecek et al., 2021). Duplicate reads were removed with the MarkDuplicates function (--REMOVE_DUPLICATES true) from the Picard package within GATK4 v4.2.5.0 (McKenna et al., 2010). Single Nucleotide Polymorphism (SNP) variants were called across autosomes with BCFtools v1.20 mpileup (-q 20 -Q 20 -Ou) and call functions (-c -Ov), and a diploid consensus sequence was created (vcfutils.pl vcf2fq -d 10 -D 100) as per the recommended methods (Li & Durbin, 2011). From this, an input file was generated for the PSMC analysis with the ‘fq2psmcfa’ function (–q 20), before population demographic history was calculated with the ‘psmc’ function (–N25 –t15 –r5 –p 4 + 25*2 + 4 + 6) from the PSMC v0.6.5-r67 package (Li & Durbin, 2011) with 100 bootstraps for each haplotype. The final PSMC plot was scaled using a tiger shark mutation rate of 1.93 × 10^−8^ (Lesturgie, Lainé, et al., 2022; Lesturgie, Planes, et al., 2022) and a generation time of 20 as the predicted range is 17.5-22.5 years (Ferreira & Simpfendorfer, 2019).

Runs of Homozygosity (ROH) were identified and used to assess the level and timescale of inbreeding occurrence as well as genes under positive selection as it is thought to be associated (Stanhope et al., 2023). Variants were again called with BCFtools mpileup (-C50 -q20 -Q20 -Ou -a DP,AD -A -f) and call (-m -A -Oz) using the same BAM files with duplicate reads subsequently removed as previously described for the PSMC analysis. The resulting VCF file was filtered with BCFtools to only retain biallelic variant sites (-m2 -M2 -v snps) and exclude variants with insufficient (1/3 genome-wide average) or excess (2x) read depth or quality (’QUAL<20 && INFO/DP<21 && INFO/DP>120’). The filtered variants were converted to PLINK format, and ROHs assessed with the homozyg function in PLINK v1.90b7.4 (Chang et al., 2015) used to define 1 Mbp (--allow-extra-chr --no-pheno --allow-no-sex --homozyg-window-het 1 --homozyg-window-missing 5 --homozyg-window-snp 50 --homozyg-density 50 -- homozyg-gap 1000 --homozyg-window-threshold 0.05 --homozyg-snp 100 --homozyg-kb 1000 -- biallelic-only) and >100kb ROH regions (--homozyg-kb 100). Associations between ROHs, CpG 5mC methylation, gene content and repeat density were tested within each chromosome using Spearman’s rank correlations across 100 kb windows, with significance assessed using a circular-shift permutation test (5,000 permutations) to account for spatial autocorrelation between adjacent windows.

Genome-wide heterozygosity was estimated using the unfiltered VCF file generated for the previous step. This was refiltered to again exclude sites with insufficient or excess read depth and low quality, however only multiallelic sites were removed while invariant sites were retained (bcftools view -i ‘N_ALT<=1’). Per-site heterozygosity was calculated in non-overlapping 1-Mbp sliding windows across the genome. Heterozygosity was defined as the number of heterozygous genotypes divided by the total number of called genotypes in each window, with the denominator including both variant and invariant positions.

## 3 RESULTS

### 3.1 The tiger shark genome

PacBio HiFi and ONT ultra-long reads, scaffolded with Hi-C Illumina sequencing, resulted in two highly complete and contiguous phased haplotype genome assemblies for the tiger shark (Table 1; Figure S1). These assemblies represent the two homologous chromosome sets of the diploid genome, with haplotypes phased based on sequence variation and read-linkage information, but without assignment to maternal or paternal origin. The final haplotype 1 assembly length of 3.54 Gbp was close to the predicted total length of 3.74 Gbp (Table 1). Haplotype 1 had 97% of scaffolds assigned to chromosomes with a scaffold N50 of 96.73 Mbp, scaffold L50 of 12, and 0.008% gap rate (Table 1). BUSCO analysis showed 97% genes were complete in both haplotype assemblies, 95% of which were single copy (Figure S2). Preliminary genome annotation produced 16,023 genes, 16,023 mRNAs and 140,580 CDSs. The final assembly consisted of 42 autosomes in each haplotype and two sex chromosomes, SUPER_X (21.9 Mb) and SUPER_Y (15.7 Mb), in haplotype 1 of this male specimen (Figure 1; Figure 2.A). This was confirmed through coverage analysis of additional known male and female samples sequenced (mean depth = 35.9X), as the females displayed double the mean coverage of the X chromosome (35.7X), compared to males (17.4X), while only males had significant coverage of the proposed Y chromosome (9.97X). Concatenation-based molecular dating positioned the tiger shark (family Galeocerdonidae) diverging between the small spotted catshark (*Scyliorhinus canicula*; family Scyliorhinidae) and the great hammerhead (*Sphyrna mokarran*) and grey reef shark (*Carcharhinus amblyrhynchos*) (Figure 2. B). This aligns with the hammerhead (Sphyrnidae) and requiem shark families (Carcharhinidae) forming one clade, with Galeocerdonidae separate within the Carcharhiniform order.

**Figure 1.**
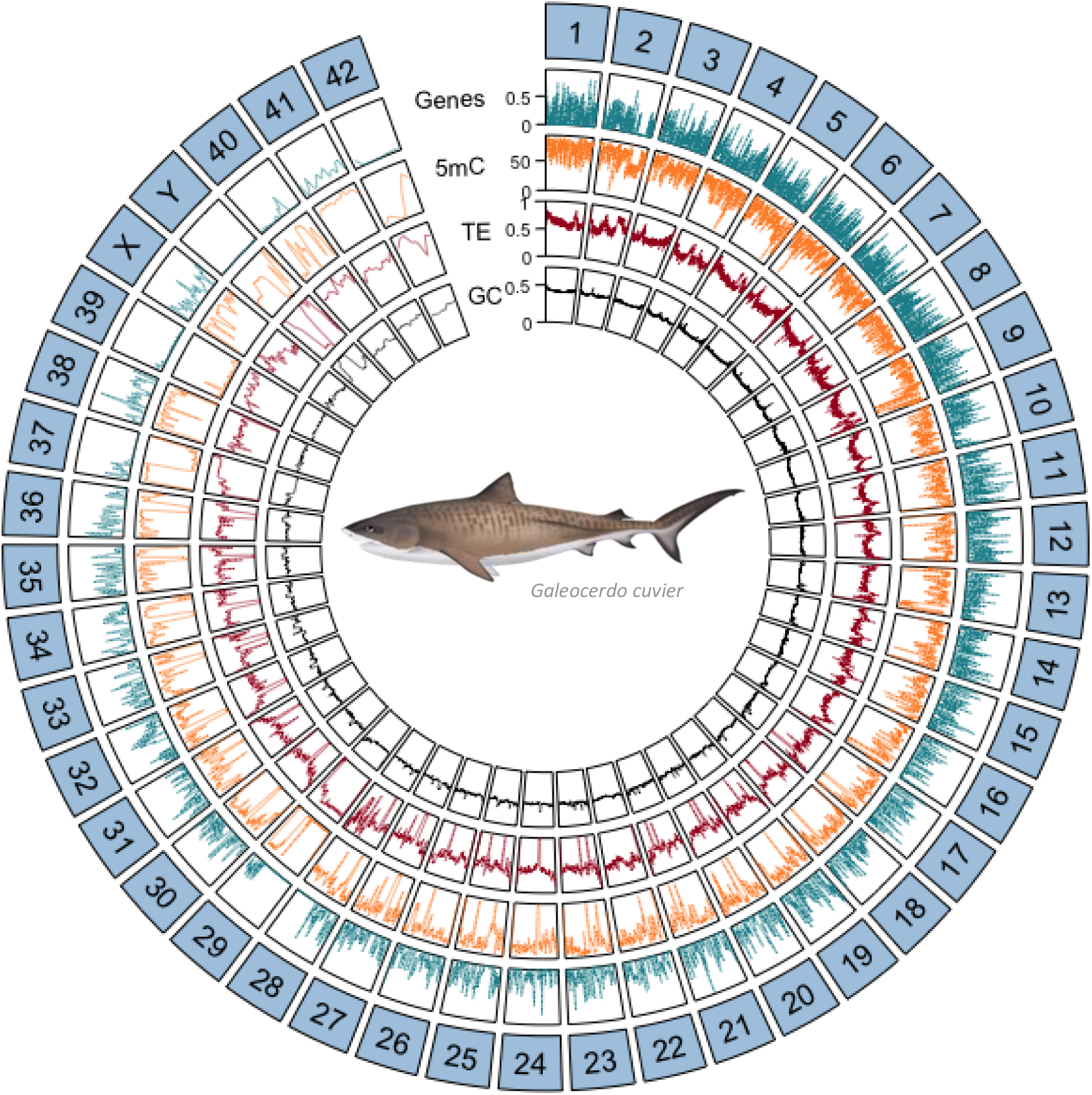
Genomic characteristics of the phased haplotype 1 chromosomes of *Galeocerdo cuvier* sGalCuv2 reference assembly. From outer to inner tracks: chromosomes (numbered and ordered by length), gene density, methylation percentage (5mC), repeat density (TE) and GC content. All tracks were calculated per 500Kbp window. Fish graphic © Marinewise.com.au

**Figure 2.**
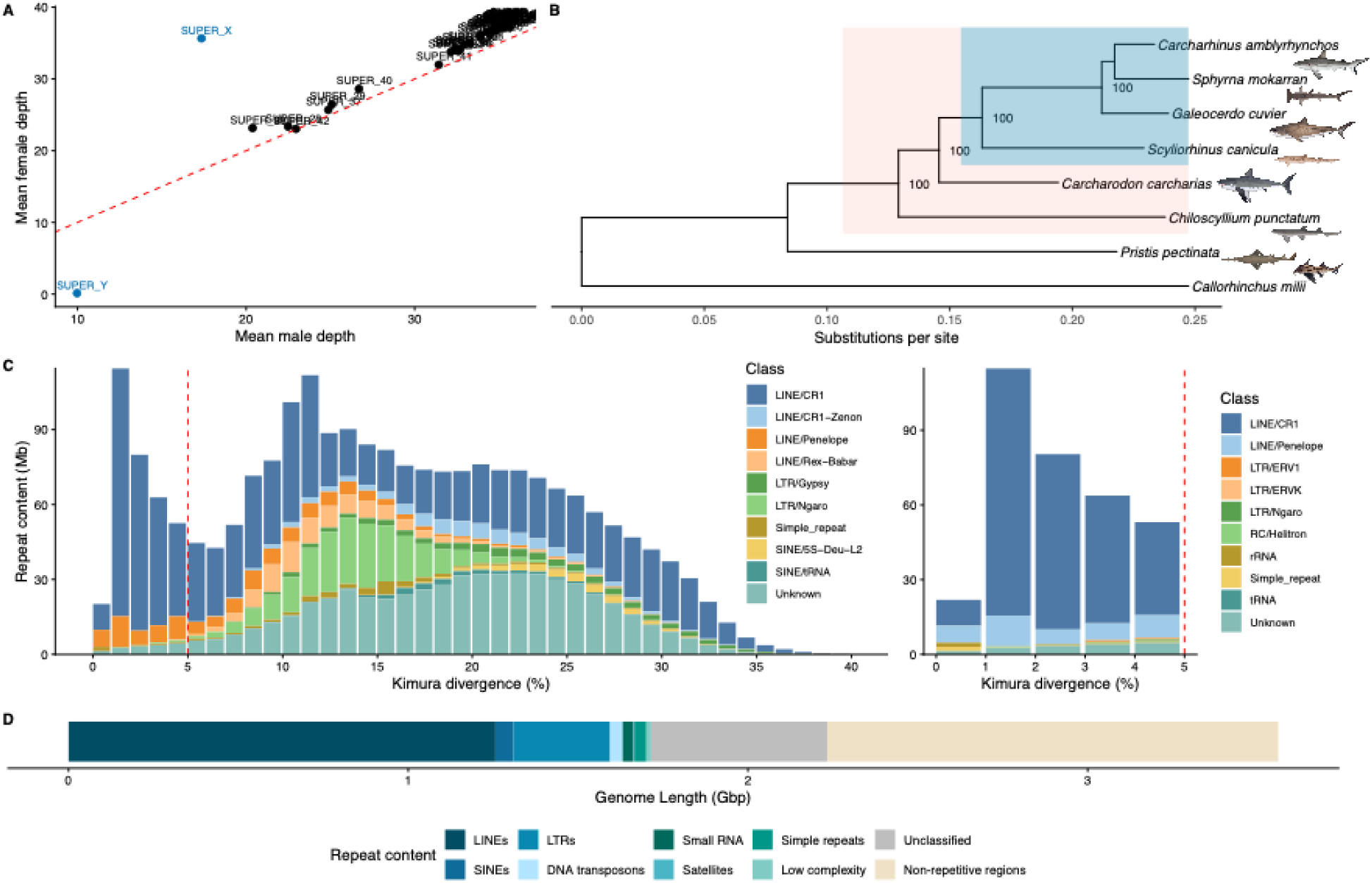
Genomic characteristics of the phased haplotype 1 chromosomes of *Galeocerdo cuvier* sGalCuv2 reference assembly continued. A) Mean coverage per chromosome (SUPER) compared between male and female samples. Red line dictates the 1:1 ratio, with sex chromosomes as outliers in blue. B) Phylogenetic tree with concatenation-based molecular dating and fossil calibration from TimeTree. Clades Selachii (red) and Carcharhiniformes (blue) are highlighted. Fish graphics © Marinewise.com.au C) Kimura divergence of repeat content by class. Left: Full Kimura divergence spectrum across whole sGalCuv2 genome. Right: Low divergence repeat content (<5%). D) Total repeat content by class.

**Table 1.** Phased genome assembly summary statistics for *Galeocerdo cuvier* (tiger shark) sGalCuv2. Complete Benchmarking Universal Single-copy Orthologs (BUSCO) ‘S’ denotes single copy and ‘D’ denotes duplicate. Merged columns reflect combined (i.e., haplotype 1 and haplotype 2) assembly statistics.

| **Metrix** | **VGP\* standards<br>(7.c.P6.Q50.C95)** | ***Galeocerdo cuvier*** |  |
|---|---|---|---|
|  |  | **Haplotype 1** | **Haplotype 2** |
| Total length (Mb) | - | 3,541.05 | 3,627.61 |
| Number of contigs | - | 1,965 | 2,129 |
| Number of scaffolds | - | 613 | 712 |
| Contig NG50 (Mb) | >10 | 5.04 | 4.88 |
| Scaffold NG50 (Mb) | = Chr. NG50 | 93.40 | 96.32 |
| BUSCO complete (%) | >95 | 96.8% [S:94.8%, D:2.0%] | 97.0% [S:94.9%, D:2.1%] |
| Gaps | < 200 per Gb | 381/Gb | 390/Gb |
| Sequences assigned to chromosomes (%) | > 95 | 96.63 | 93.95 |
| QV | >50 | 60.66 | 59.48 |
|  |  | 60.02 |  |
| Merqury completeness (%) | >95 | 96.12 | 95.95 |
|  |  | 99.47 |  |

Characterising the non-coding landscape of this genome assembly revealed tiger sharks display high amounts of repetitive content, accounting for 62.51% of the total genome length (Figure 2.D; Table S3). Retroelements made up 45%, with Long interspersed nuclear elements (LINEs) retrotransposons being the largest class of repetitive element at 35.5% (1.26Gbp), almost all of which was L2/CR1/Rex type elements (29.3%; 1.04Gbp). Unclassified interspersed repeats were the next highest class (14.6%), followed by Long Terminal Repeat (LTR) retrotransposon elements (8%). Repeat content was often enriched at the ends of chromosomes around the telomeres (Figure 1).

In the comparative analyses, the tiger shark displayed the largest proportion of recent repeat expansion (≤5% Kimura divergence), comprising 9.65% of the genome (Figure 2.D; Table S2). This was predominantly driven by a major recent expansion of LINE/CR1 elements (7.69%), with additional contributions from Penelope, LTR and unknown transposable elements (Figure 2.D; Table S2). Simple repeats also showed substantial recent expansion (Figure 2.D; Table S2). This pattern in the repeat divergence landscape was clade specific, as the great hammerhead (*Sphyrna mokarran*) and grey reef sharks (*Carcharhinus amblyrhynchos*) also displayed significant recent expansions in LINE/CR1, Penelope and unknown transposable elements (Figure S4), as well as a higher proportion of recent divergence in general (7.35% and 6.89% respectively). The small spotted catshark (*Scyliorhinus canicula*) also displayed a higher proportion of recent expansions (8.46%) and while LINE/CR1 elements remained predominant in this, LTR/Ngaro and unknown elements became equally as prominent (Figure S4).

Repeat Kimura divergence landscape patterns were vastly different between orders (Figure S4). Outside of Carcharhiniformes, but within the elasmobranch subclass, the white shark (*Carcharodon carcharias*), brownbanded bamboo shark (*Chiloscyllium punctatum*) and smalltooth sawfish (*Pristis pectinata*) displayed decreased proportions of recently diverged repeat content (2.92%, 4.96% and 2.93% respectively) and all retained substantial proportions of recent LINE/CR1, simple and unknown element expansions. Interestingly, recent repeat expansion in the white shark was largely driven by LINE/Penelope elements, but LINE/Rex-Baber and LTR/Gypsy in the sawfish. The elephant shark (*Callorhinchus milii*) unsurprisingly displayed the most different repeat divergence landscape (Figure S4) as the only chimaera included in this analysis. Total recent content was 3.7% and characterised by SINE/ID and LINE/L2 elements, although also displayed substantial proportions of recent simple and unknown elements (Figure S4).

### 3.2 The tiger shark methylome

We used PacBio HiFi and ONT ultra-long read sequencing to characterise methylation patterns across the genome. Global mean 5mC methylation was 62.88% across CpG sites with at least 10X PacBio HiFi coverage (Table S5). ONT data was lower coverage overall (mean sequencing depth = 8.8X), and so inference was limited to sites with a minimum of 5X. An average CpG 5mC methylation of 63.42% was maintained at this lower threshold (Table S5). As a 5X filtering threshold closely matched the results of the high coverage HiFi data, this was deemed adequate for all ONT derived methylation estimates. The methylation distribution was bimodal for both HiFi and ONT derived CpG 5mC, but skewed towards higher methylation frequencies, with most sites exhibiting either near zero or above 60% methylation (Figure 3.A; Figure S5). Almost 75% of gene body regions averaged over 80% methylation, whereas close to 50% of promoter regions had an average under 20% (Figure 3B). This pattern was observed for both HiFi and ONT CpG 5mC derived estimates (Figure 3.B; Figure S6). CpG 5mC methylation was largely consistent across major chromosomes, with the noticeable exception of chromosomes 29, 37 and 39, which displayed extreme hypomethylation (Figure 3C; Figure S7.A). These regions appear to correlate with extremely low gene content and particularly high repeat density (Figure 2; Figure 3C). However, only CpG 5mC methylation and gene content were significantly correlated for chromosome 29 (Spearman’s rank = 0.55, p = 0.002) and chromosome 37 (Spearman’s rank = 0.44, p < 0.001), while neither were significant for chromosome 39.

**Figure 3.**
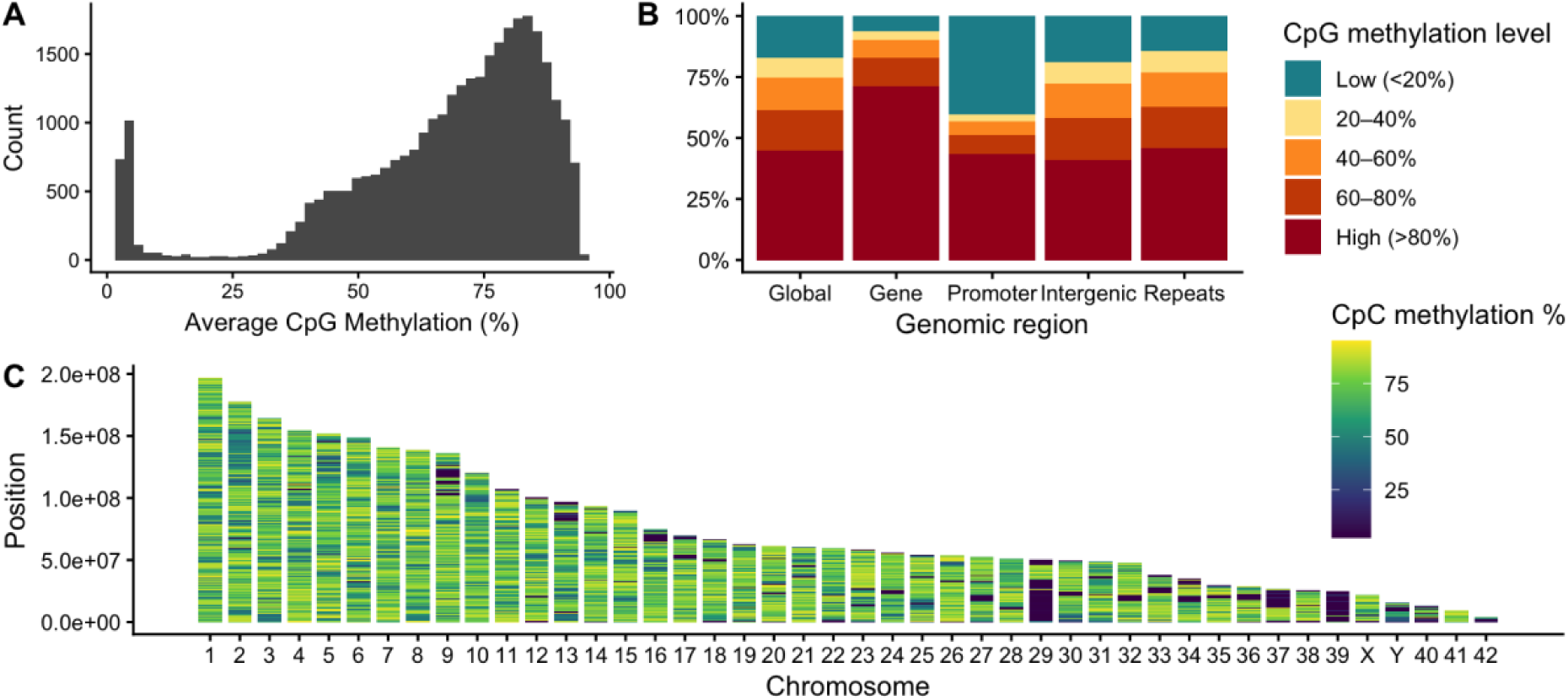
Genome wide methylation profile for the tiger shark (*Galeocerdo cuvier*) sGalCuv2 reference assembly called from PacBio HiFi data. A) Methylation distribution as the average percentage of CpG sites across 100KB windows B) Average methylation level for various genomic regions. C) Average CpG methylation level for 100KB windows across all chromosomes ordered by length.

ONT data also has the ability to call 5mC methylation across all cytosines (all-context), as well as 5hmC methylation for both CpG sites and all cytosines. Global average all-context 5mC methylation was predictably much lower at only 2.6% of all filtered sites (Table S5), and displayed a unimodal distribution, with most of the genome under 5% (Figure S5). A similar pattern across genomic regions was observed nonetheless, with gene regions being the most methylated and promoter regions the least, but at a considerably lower level (Figure S6). Chromosome 28 however displayed an elevated level of all-context 5mC methylation (Figure S7). Conversely, 5hmC methylation averaged 4.07% within CpG sites, and 0.08% across all cytosines (Table S5). Methylation distribution was unimodal for both, with all CpG sites under an average of 10% 5hmC methylation and under 0.25% across all cytosines (Figure S8). There were minimal differences in methylation level between all genomic regions investigated for either (Figure S9), although once again gene bodies and promoter regions exhibited slightly elevated and lower levels respectively within CpG sites. Chromosomes 28 again presented higher levels of 5hmC all-context methylation, along with the X and Y chromosomes (Figure S10).

### 3.3 Demographic history across deep time

Genome-wide heterozygosity provides a measure of long-term genetic processes, while the distribution and lengths of runs of homozygosity (ROH) provide complementary insight into the timing and extent of demographic change and inbreeding. Together they reveal historical population dynamics and their potential consequences for contemporary genetic resilience. Genome-wide heterozygosity for the tiger shark was low at 1.35/kb, whilst ROHs were found throughout genome (Figure 4). Heterozygosity increased towards the distal edges of larger chromosomes and was elevated on average in smaller ones (Figure 4.C). The length and number of ROH reflect levels of individual inbreeding, with longer ROH indicative of recent inbreeding and shorter ROH suggestive of more historical events due to recombination events. Longer ROHs (>1 Mbp) totalled almost 30 Mbp between 21 segments across the 42 autosomes of the tiger shark genome. These segments averaged 1.42 Mbp in length but ranged from 1 to 2.4 Mbp in size and translated to a low inbreeding coefficient of FROH_(>1Mbp)_ = 0.9%. Although small in number, ROHs > 1 Mbp were detected throughout the genome, including large chromosomes 2 to 9, to smaller chromosomes 31 to 36 (Figure 4.D). Strikingly however, 43% of segments were identified within chromosome 28. Smaller ROHs (>100kbp) had a higher inbreeding coefficient (FROH_>100kbp_) with 7.2% of the genome in ROH (Figure 4.D). These totalled 233Mbp across 1034 segments, averaging 225Kbp in size. Haplotype 2 displayed similar, albeit slightly higher, values across all ROHs investigated. Regions of extremely low heterozygosity appeared to correlate with the longest and most dense ROHs, most noticeably on chromosomes 28 and 29, as well as lower gene content, high repeat density and hypomethylation. ROHs did significantly correlate with decreased CpG 5mC methylation and increased repeat density for chromosomes 2 (CpG 5mC: Spearman’s rank = -0.2, p = 0.049; Repeat density: Spearman’s rank = 0.28, p = 0.011). However, no other factors, except heterozygosity, were significant for chromosomes 28, 29, 37 and 39 where these runs were apparent.

**Figure 4.**
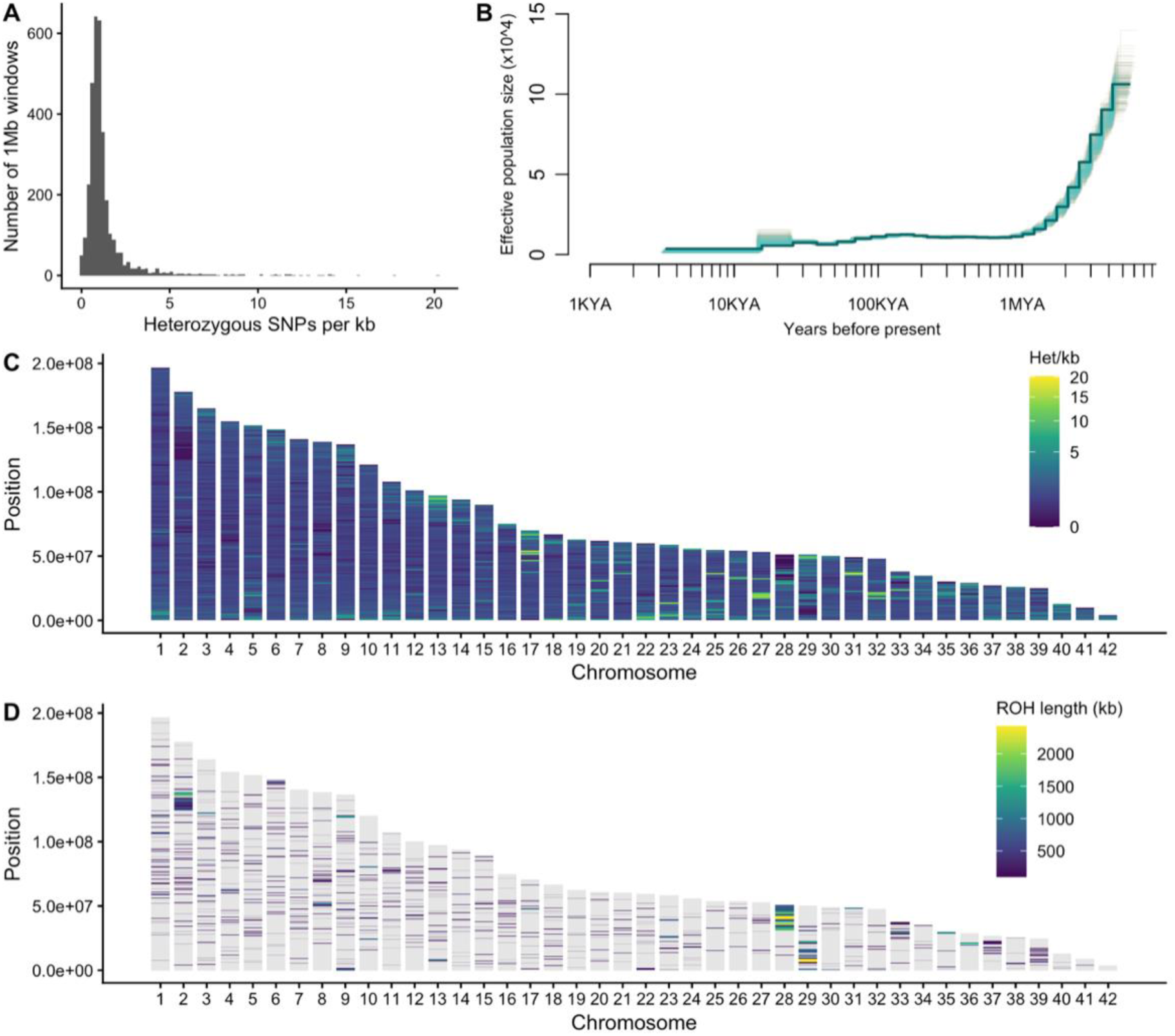
Patterns of demographic history in the tiger shark. A) Distribution of heterozygosity per kb in 1Mb windows B) Effective population size (N_e_) of tiger sharks across the last 8 million years, calculated using a PSMC model with 100 bootstrap replicates for both Haplotypes 1 (blue) and 2 (orange). C) Average heterozygosity per kb and D) Runs of Homozygosity (ROH) >100Kbp both calculated in non-overlapping 100kb windows across all chromosomes ordered by length.

Reconstruction of tiger shark historical effective population size (N_e_) through a PSMC model was also consistent between haplotypes, revealing a gradual decline from an N_e_ of roughly 100,000 to 20,000 between 8 million years ago (MYA) and 1 MYA (Figure 4.B). Following this, N_e_ plateaued temporarily until a further decline to 10,000 after 100 KYA. Noticeably, bootstrapping variability increases dramatically during the period between 15 to 30 KYA, before a sharper decline in N_e_ occurs (Figure 4). After this final drop to 1,000 around 15 KYA, N_e_ once again plateaued until the limit of this model at 3,000 years ago.

## 4 DISCUSSION

In this study, we utilised multiple sequencing platforms to produce a highly complete and contiguous phased haplotype genome assembly for the tiger shark. Our results provide new insights into genomic and epigenomic characteristics, and demographic history of this species. The chromosome-scale assembly identified an X and putative Y sex chromosomes, while comparative analyses identified lineage-specific repeat evolution characterised by a pronounced recent CR1 expansion. Genome-wide methylation analyses further revealed distinct patterns of CpG and all-context methylation and provided the first 5hmC methylome sequencing of a chondrichthyan. Finally, low genome-wide heterozygosity, runs of homozygosity, and a prolonged decline in effective population size point to sustained demographic contraction rather than a single severe bottleneck. Together, these findings demonstrate the complex evolutionary history of the tiger shark and establish a genomic framework for understanding its genetic resilience and vulnerability under contemporary environmental change.

The final assembly matched the known (2n = 86) karyotype in terms of numbers of chromosomes, but heterogamety was previously indeterminate based on the karyotype, despite XY appearing as the dominant sex system in chondrichthyan species (Maddock & Schwartz, 1996; Niwa et al., 2025; Schwartz & Maddock, 2002). Thus, this study presented the first evidence of XY sex determination in this species. We demonstrated diploid coverage of the X chromosome and near zero of the Y in female samples, with half coverage of both in males consistent with a hemizygous state (Wu et al., 2024; Yamaguchi et al., 2023). The Y chromosome is typically extensively degenerated, around one-third the size of the X, across cytogenetic and genome assembly identification of XY systems in sharks (Dahms et al., 2025; Uno et al., 2020; Wu et al., 2024). However, the tiger shark Y chromosome presented here is two-thirds the size of the larger X and we could not identify a pseudoautosomal region (PAR). The notably high GC content across one section suggests there is excessive secondary or non-canonical structure occurring within the Y chromosome that could result in poor sequencing and could be falsely inflating the size. The lack of a PAR region also could be due to collapsed representation or incomplete assembly of the Y chromosome. BUSCO completeness (95%) for this assembly was standard, if not higher, than many shark genomes (Dahms et al., 2025; de Jong et al., 2026; Yamaguchi et al., 2023), yet genome annotation yielded a low level of genes compared to other sharks with transcriptome evidence (Wu et al., 2024), although it was similar to those without (Dahms et al., 2025). Consequently, this chromosome assembly should not currently be considered complete and identification of it as the putative Y is provisional (Niwa et al., 2025).

Comparative repeat landscapes indicated that recent transposable element activity has been lineage-specific among sharks. The tiger shark genome displayed high repeat content like that of other closely related shark species (Dahms et al., 2025; Laso-Jadart et al., 2025; Stanhope et al., 2023). Expansions of repeat content, particularly of CR1-like LINES, are prolific across shark species. These elements contribute to generally large genome sizes but are also associated with fragile chromosome sites prone to breakages (Weber et al., 2020; Wu et al., 2024). Tiger shark also displayed the highest proportion of recently expanded repeat content in our comparative analyses. The pronounced recent CR1 expansion across Carcharhiniformes suggests ongoing or recent retrotransposon activity that may have contributed to genome restructuring and regulatory evolution in this lineage. The large proportion of unknown repeat elements contributing to recent expansions could also indicate poorly classified class specific TEs. Rather than a conserved pattern of repeat expansion, batoids and chimeras exhibited amplification of distinct TE families, suggesting independent episodes of TE activation following species divergence.

### 4.1 Conserved and chondrichthyan-specific patterns revealed by the tiger shark methylome

The close agreement between PacBio HiFi and ONT estimates of global and feature-level CpG methylation provides independent support for the broad methylation patterns observed, despite differences in sequencing depth and modification-calling approaches. Consistent with the hypermethylated genomes characteristic of most vertebrates (Suzuki & Bird, 2008), global CpG methylation level in the tiger shark was 63%, marginally lower than the 69–72% previously reported in other chondrichthyans including the elephant shark (Peat et al., 2017) and school shark (de Jong et al., 2026). The pronounced contrast between hypermethylated gene bodies and predominantly hypomethylated promoters demonstrates strong functional partitioning of CpG methylation across the tiger shark genome. This organisation is characteristic of vertebrate methylomes, where promoter hypomethylation generally maintains regulatory regions in a transcriptionally permissive state, while gene-body methylation has been associated with transcriptional regulation and suppression. Hypermethylation was similarly displayed across >50% repetitive elements, conforming to the pattern of other chondrichthyans and vertebrates (Peat et al., 2017) and the established role of DNA methylation in transposable element silencing.

Interestingly, chromosomes 29, 37 and 39 displayed extensive hypomethylated regions that coincided with low gene content and high repeat density (Figures 2 and 3C), although statistical support was limited. Gene content was significantly correlated with CpG 5mC methylation on chromosomes 29 and 37, but not 39. Although repetitive sequences are typically heavily methylated, hypomethylation of repetitive DNA is not without precedent, such as satellite repeat arrays in the human genome (Hoyt et al., 2022). Gene-poor, repeat-rich microchromosomes have been shown previously in shark species (Wu et al., 2024) and can carry distinct epigenetic landscapes, as has been reported in the maize B chromosome (Liu et al., 2025). While hypomethylation has previously been associated with high SNP density in sharks (de Jong et al., 2026), the regions identified here instead overlapped with ROHs and areas of low heterozygosity (Figures 3 and 4). This was most pronounced on chromosomes 28 and 29, where long, dense ROHs overlapped with reduced gene content and high repeat density, alongside lower methylation on chromosome 28 (∼50%) and extreme hypomethylation on chromosome 29. However, statistical associations between ROHs and reduced CpG 5mC methylation and gene content or increased repeat density were again limited, with significant relationships detected only on chromosome 2. Collectively, these patterns suggest that regional methylation may be influenced by underlying genomic architecture, including gene and repeat content, and not necessarily localised genetic diversity.

Although DNA methylation in vertebrates occurs predominantly in the CpG context, non-CpG methylation has also been implicated in gene expression, transposable element regulation and hence genome stability as well, but the extent to which remains uncertain (Deniz et al., 2019). Most cytosines are not in CpG context and subsequently all-context methylation is generally low, around <1–2% in vertebrates (He & Ecker, 2015; Lister et al., 2009). The tiger shark exhibited over double the levels of all-context 5mC methylation (2.6%; Figure S5 and S7) than the elephant shark (*Callorhinchus milii*; 0.8– 1%), the only other chondrichthyan species with published estimates (Peat et al., 2017). However, given that these estimates are currently available for only two distantly related chondrichthyan species, it is not possible to determine whether this difference is an artifact of whole genome bisulfite sequencing verses ONT or genuinely elevated non-CpG methylation in the tiger shark.

Hydroxymethylation of CpGs (5hmC methylation) was initially thought to be purely an intermediate in DNA demethylation but is now associated with transcriptionally permissive chromatin (Pastor et al., 2013; Wu & Zhang, 2017). However, very little research into 5hmC methylation has been done outside of mammals and has only been reported for one chondrichthyan species before using mass spectrometry (global 5hmC of 0.01%) and pyrosequencing of specific CpG sites (Colwell et al., 2018). Hydroxymethylation was detected at 4.07% of CpG sites and 0.08% across all cytosines throughout the tiger shark genome (Figures S8 and S10). The all-context 5hmC level was comparable to global 5hmC levels reported for non-mammalian vertebrates (∼0.075%) (Colwell et al., 2018; Pastor et al., 2013; Wu & Zhang, 2017), but potentially indicating CpG enrichment of 5hmC in sharks. As 5hmC functions as both an intermediate in active DNA demethylation and a regulatory epigenetic mark, its presence indicates that dynamic methylation turnover is maintained in chondrichthyan genomes. The exact extent however should be interpreted with caution as the modification-calling model has not been validated in a non-model organism.

### 4.2 Tiger sharks display signals of long-term low N_e_ and heterozygosity

The PSMC reconstruction indicated a gradual historical decline in effective population size (N_e_) beginning approximately 8 million years ago and continuing until around 1 MYA, followed by relative stability through much of the Pleistocene (Figure 4B). This period of relative stability followed substantial turnover within the tiger shark lineage (genus: Galeocerdo) during the late Miocene to Pliocene, when several extinct species, including *G. aduncus*, *G. mayumbensis* and later *G. capellinii*, disappeared, leaving *G. cuvier* as the sole surviving representative by approximately 2–5 MYA (Türtscher et al., 2021). Although ecological competition may have contributed to the disappearance of these congeners, given their potentially overlapping trophic roles, this remains speculative (Höltke et al., 2024; Türtscher et al., 2021). The PSMC trajectory differed from previous demographic studies of Indo-Pacific tiger sharks, which inferred an expansion around 100,000–200,000 years ago, followed by a substantial decline or bottleneck approximately 1,600–4,000 years ago (Lesturgie, Lainé, et al., 2022; Lesturgie, Planes, et al., 2022). In contrast, our reconstruction indicated a further decline from approximately 100,000 to 15,000 years ago, followed by persistently low N_e_, without clearly resolving the more recent bottleneck. These differences may partly reflect methodological differences, as PSMC reconstructs coalescent history from a single diploid genome, while Lesturgie et al. inferred demographic changes from the site frequency spectrum across multiple individuals, providing greater resolution of recent demographic events. Although our analysis is based on a single genome from one sampling location, previous studies found weak to negligible population structure across the Indo-Pacific, suggesting that population structure is unlikely to fully explain these differences. However, the generation time used in this study was based on the most recent IUCN assessment (∼20 years) and was double that from Lesturgie et al. (10 years), which would have affected only the estimated timing of demographic events (Robinson et al., 2021). Overall, these findings highlight the complexity of reconstructing tiger shark demographic history and the need for broader whole-genome sampling and complementary demographic approaches to better resolve historical population changes (Lesturgie, Lainé, et al., 2022).

This genome also presented evidence of historical, and potentially recent, inbreeding in tiger shark populations. Heterozygosity was 1.35/kb, with a moderate proportion of short ROHs (FROH>100Kbp) and a small fraction of long ROHs (FROH>1Mbp). Heterozygosity observed in this tiger shark was larger than that observed for other large-bodied, wide-ranging shark species such as the white shark (*Carcharodon carcharias*; 0.97/kb) and whale shark (*Rhincodon typus*; 0.65/kb) (Stanhope et al., 2023). Yet, this was considerably lower average heterozygosity than that of the shortfin mako (*Isurus oxyrinchus*; 3.58/kb), despite half the proportion of short ROHs and a similar level of long ROHs. The shortfin mako historically had a high N_e_, but underwent an immense population crash around 100KYA (Stanhope et al., 2023). Based on the PSMC, the tiger shark did not experience such a severe bottleneck at one time, but a series of smaller declines, resulting in fewer ROHs. We hypothesise that sustained lower N_e_ over a longer time instead produced the lower heterozygosity observed in the tiger shark. Inbreeding resulting in ROHs over 100kb detected likely occurred around a few thousand years ago, matching the PSMC timeframe (Stanhope et al., 2023). Few long ROHs over 1Mb implied relatively little contemporary inbreeding in the past 200 years (Stanhope et al., 2023). However, all methods for detecting ROHs tend to underestimate FROH and are sensitive to many demographic and data variables (Silva et al., 2024), suggesting these estimates could be higher in reality.

This low genetic diversity seems to be characteristic for numerous shark species (Feutry et al., 2014; Stanhope et al., 2023), potentially as elasmobranchs exhibit reduced molecular evolutionary rates relative to many other vertebrates (Hara et al., 2018; Martin et al., 1992; Sendell-Price et al., 2023). While this could indicate reduced capacity for future adaptation, sustained low N_e_ and heterozygosity may not necessarily have deleterious impact (Domingues et al., 2018; Hohenlohe et al., 2021). The long ROHs shown in the tiger shark on chromosomes 28 and 29, could also reflect regions undergoing selection or associated with adaption resulting in a lack of recombination (Biscarini et al., 2026). Regardless, this study demonstrates that tiger shark populations are sensitive to climate shifts which could be compounded by contemporary anthropogenic pressure.

### 4.3 Reference genomes are foundational for high-resolution conservation inference

The reference genome and associated genomic resources developed here provide an important foundation for high-resolution conservation research in tiger sharks, although several limitations remain. The current genome annotation should be considered provisional, with future RNA-seq-based annotation needed to improve gene predictions and functional interpretation. Similarly, the methylation patterns presented here represent blood from a single male sampled at one location; broader sampling across tissues, sexes, geographic regions and environmental conditions will be necessary to characterise epigenetic variation and its potential regulatory significance in chondrichthyans. The PSMC demographic reconstruction also requires cautious interpretation, as N_e_ estimates and the timing of demographic events are sensitive to mutation-rate and generation-time assumptions (Robinson et al., 2021). Although the reconstruction extends to approximately 8 MYA, fossil evidence suggests that the extant tiger shark lineage originated around 13.8 MYA (Türtscher et al., 2021), meaning that earlier evolutionary history is not captured by this analysis. Furthermore, PSMC has limited resolution of recent demographic changes, particularly over the past millennium, and contemporary estimates of tiger shark N_e_ remain unavailable. Future whole-genome studies incorporating broader geographic and temporal sampling will therefore be important for establishing contemporary demographic baselines and monitoring population trends.

Although still relatively few cartilaginous fish have available reference genome assemblies, dedicated projects in this space are now working to remedy this given the high ecological, social, and economic importance of this clade (Nishimura et al., 2022; Parata, de Jong, et al., 2025). This study is an essential step in the investigation of chondrichthyan genomes at the chromosome scale and in understanding their evolution. Given genetic differentiation between the Indo-Pacific and the Atlantic tiger shark populations (Bernard et al., 2021), production of an Atlantic reference assembly would be beneficial and an interesting point of comparison. The reference genome presented here provides a stepping stone for future understanding into tiger shark evolution and historical population dynamics, as well as insight into a unique evolutionary path across all sharks. Nevertheless, the true power of a reference genome lies in its integration with population genomic data, enabling contemporary patterns of adaptation and connectivity to be resolved in the tiger shark at unprecedented genomic resolution.

## Data and code availability

The tiger shark sGalCuv2 genome assembly has been deposited at DDBJ/ENA/GenBank under the accessions JCFYQQ000000000 (Haplotype 1) and JCFYQR000000000 (Haplotype 2). The versions described in this paper are JCFYQQ010000000 and JCFYQR010000000. The preliminary genome annotation is available on FigShare under DOI: 10.6084/m9.figshare.34882005. Raw sequence data and assemblies for the additional four high-coverage Illumina short-read individuals are available on ENA under BioSamples (assembly accessions): SAMEA123517613 (GCA_988294775.1), SAMEA123517612 (GCA_988294875.1), SAMEA123517614 (GCA_988294915.1), SAMEA123517615 (GCA_988294835.1).

## Acknowledgements

We would like to acknowledge that the University of Western Australia is situated on the land of the Whadjuk people of the Noongar nation and acknowledge their continuing connection to land and sea country. We would like to acknowledge that the samples used in this study were collected from Exmouth Gulf of the West Thalanyi-Jinigudira/Yinigurdira and Baiyungu peoples. We acknowledge collaborative support of Minderoo Foundation, the Minderoo Exmouth Research Laboratory and Minderoo OceanOmics Centre at UWA in this work and its mission to protect and restore natural ecosystems for future generations. Shark images incorporated throughout the manuscript are © marinewise.com.au.

## Funding statement

Minderoo Foundation, the Minderoo Exmouth Research Laboratory and Minderoo OceanOmics Centre at UWA funded sample collection and data generation. Additional funding support for sampling was provided by Holsworth Wildlife Research Endowment.

## Competing Interests

The authors declare no competing interests.

## Notes

### Competing Interest Statement

The authors have declared no competing interest.

