## Supplementary materials for "The tiger shark, *Galeocerdo cuvier* (Péron & Lesueur, 1822), genome reveals genomic and epigenomic signatures of shark evolution and demographic history"

### Section 1: Genomics methods

**Table S1.** Reference assembly information used for comparative genomic analyses (Class: Chondrichthyes)

| **Taxonomy** | **Species name** | | **NCBI accession** | **Citation** |
|---|---|---|---|---|
| **Subclass** | Scientific | Common | | |
| **Holocephali** | *Callorhinchus milii* | Elephant shark | GCF_018977255.1 | (Venkatesh et al., 2014) |
| **Elasmobranchii** | | | | |
| Batoidea | *Pristis pectinata* | Smalltooth sawfish | GCF_009764475.1 | (Rhie et al., 2021) |
| Selachii | | | | |
| Orectolobiformes | *Chiloscyllium punctatum* | Brownbanded bamboo shark | GCF_047496795.1 | (Niwa et al., 2025) |
| Lamniformes | *Carcharodon carcharias* | White shark | GCF_017639515.1 | (Laso-Jadart et al., 2025) |
| Carcharhiniformes | | | | |
| Scyliorhinidae | *Scyliorhinus canicula* | Smaller spotted catshark | GCF_902713615.2 | (Hara et al., 2018) |
| Carcharhinidae | *Carcharhinus amblyrhynchos* | Grey reef shark | GCF_965287045.1 | (Dahms et al., 2025) |
| Sphyrnidae | *Sphyrna mokarran* | Great hammerhead shark | GCA_024679065.1 | (Stanhope et al., 2023) |
| Galeocerdontidae | *Galeocerdo cuvier* | Tiger shark | PRJNA1245416 | This paper |

**Table S2.** Calibration points from TimeTree used for phylogenetic tree dating.

| **Taxon 1** | **Taxon 2** | **Divergence times** |
|---|---|---|
| Holocephali | Elasmobranchii | 338.0 - 471.0 MYA |
| Selachii | Batoidea | 219.7 - 343.0 MYA |
| Orectolobiformes | Carcharhiniformes | 112.9 - 289.0 MYA |
| Orectolobiformes | Lamniformes | 112.9 - 289.0 MYA |
| Lamniformes | Carcharhiniformes | 168.2 - 259.0 MYA |
| **Node** | **Split** | **100 Myr units** |
| Root | Holocephali/Elasmobranchii | B(3.38, 4.71) |
| Elasmobranchii | Batoidea/Selachii | B(2.197, 3.43) |
| Selachii | Orectolobiformes/rest | B(1.129, 2.89) |
| Lamniformes/Carcharhiniformes | Lamniformes +Carcharhiniformes | B(1.682, 2.59) |

### Section 2: Genome assembly and quality assessment

A)

B)

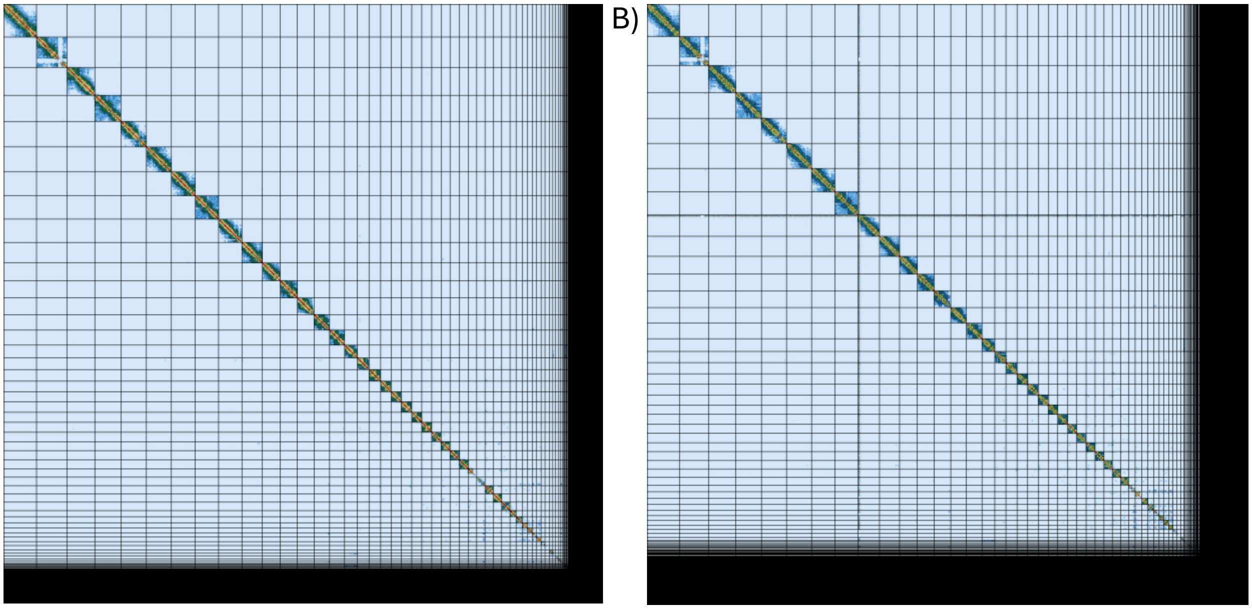

**Figure S1.** Hi-C linked contact maps of the final curated genome assembly of a) *Galeocerdo cuvier* (tiger shark) sGalCuv2 haplotype 1 and b) *Galeocerdo cuvier* sGalCuv2 haplotype 2. All chromosomes are ordered by length.

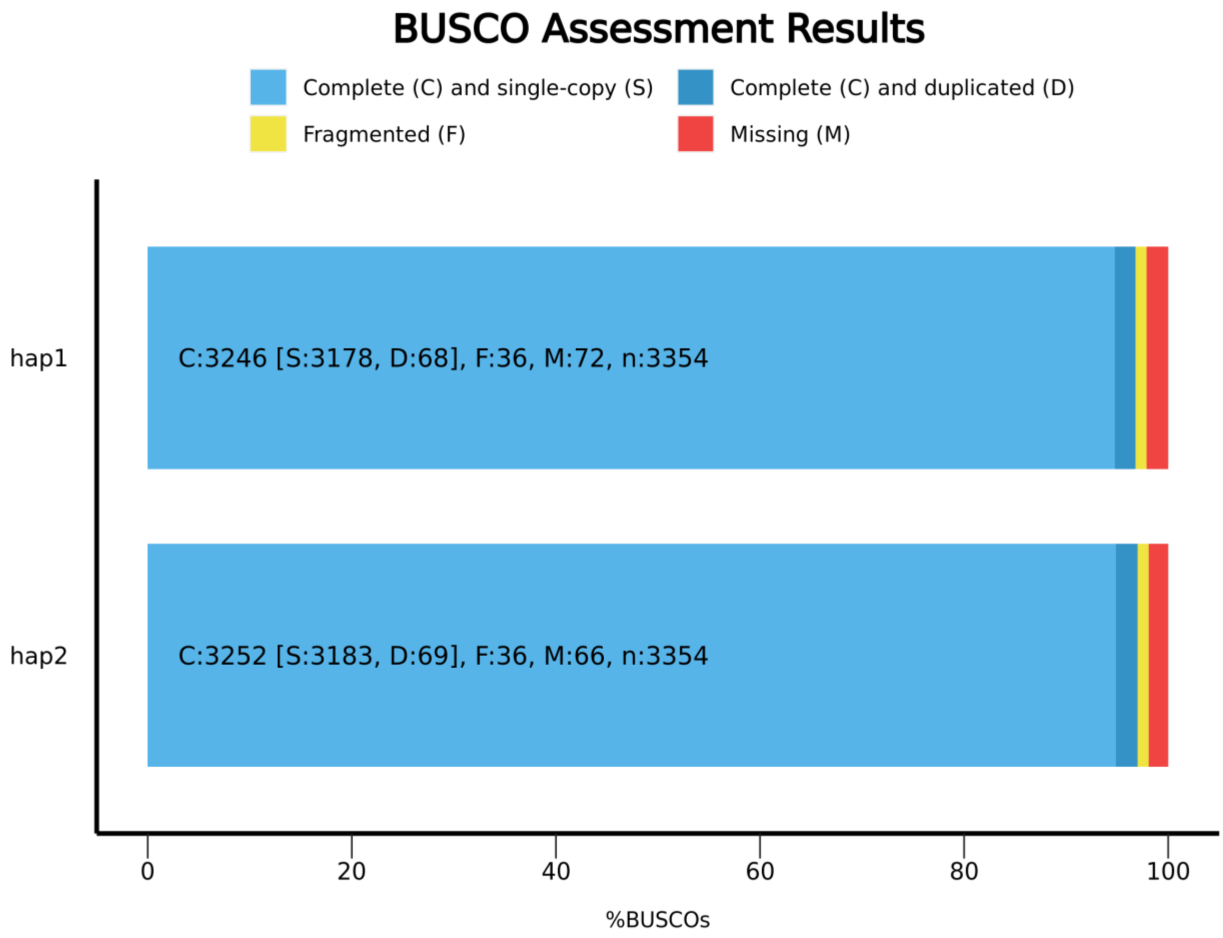

**Figure S2.** Benchmarking Universal Single-copy Orthologs (BUSCO) assessment results for *Galeocerdo cuvier* (tiger shark) sGalCuv2 reference genome assembly.

**Table S3.** Full repeat content of the *Galeocerdo cuvier* (tiger shark) sGalCuv2 haplotype 1 assembly produced by repeatmasker v4.1.5, run with rmblastn version 2.14.1+.

| Repeat type | Number of elements | Length occupied (bp) | % of genome |
|---|---|---|---|
| **Retroelements:** | **4,145,072** | **1,594,040,890** | **45.02** |
| SINEs | 329,785 | 52,894,792 | 1.49 |
| Penelope | 1,498 | 201,461 | 0.01 |
| LINEs | 2,747,444 | 1,256,588,256 | 35.49 |
| *CRE/SLACS* | *0* | *0* | *0* |
| *L2/CR1/Rex* | *2,150,298* | *1,042,170,634* | *29.43* |
| *R1/LOA/Jockey* | *1,814* | *185,194* | *0.01* |
| *R2/R4/NeSL* | *363* | *476,123* | *0.01* |
| *RTE/Bov-B* | *50,224* | *19,515,002* | *0.55* |
| *L1/CIN4* | *13,225* | *7,005,292* | *0.20* |
| **LTR elements:** | **1,067,843** | **28,455,7842** | **8.04** |
| BEL/Pao | 0 | 0 | 0 |
| Ty1/Copia | 8,490 | 4,018,103 | 0.11 |
| Gypsy/DIRS1 | 117,859 | 59,118,854 | 1.67 |
| *Retroviral* | *24,314* | *14,377,989* | *0.41* |
| **DNA transposons:** | **174,355** | **37,266,753** | **1.05** |
| hobo-Activator | 16,042 | 3,873,174 | 0.11 |
| Tc1-IS630-Pogo | 92,367 | 12,820,536 | 0.36 |
| En-Spm | 0 | 0 | 0 |
| MULE-MuDR | 0 | 0 | 0 |
| PiggyBac | 0 | 0 | 0 |
| Tourist/Harbinger | 38,589 | 14,291,844 | 0.40 |
| Other | 0 | 0 | 0 |
| **Rolling-circles** | 13,069 | 1,803,591 | 0.05 |
| **Unclassified** | 3,041,374 | 518,449,263 | 14.64 |
| **Total interspersed repeats** | | **2,149,958,367** | **60.72** |
| **Small RNA** | 176,124 | 31,748,051 | 0.90 |
| **Satellites** | 12,158 | 2,762,891 | 0.08 |
| **Simple repeats** | 205,307 | 34,874,765 | 0.98 |
| **Low complexity** | 41,906 | 14,196,955 | 0.40 |
| **Total repetitive content** | | **2,213,556,686** | **62.51** |

**Table S4.** Recently expanded repeat content within the *Galeocerdo cuvier* (tiger shark) sGalCuv2 haplotype 1 assembly (≤5% Kimura divergence).

| Class | Bases covered | Number of elements | Percentage of genome |
|---|---|---|---|
| LINE/CR1 | 272156001 | 237399 | 7.685742958 |
| LINE/Penelope | 41533365 | 81895 | 1.172911001 |
| Unknown | 15656799 | 55977 | 0.442151311 |
| Simple_repeat | 3189027 | 68177 | 0.090058796 |
| rRNA | 1713048 | 348 | 0.048376837 |
| LTR/Ngaro | 1543260 | 13970 | 0.043581988 |
| LTR/ERV1 | 1259325 | 601 | 0.035563604 |
| tRNA | 702797 | 4671 | 0.019847136 |
| LTR/ERVK | 642254 | 2460 | 0.018137389 |
| RC/Helitron | 637314 | 1245 | 0.017997882 |
| LINE/Rex-Babar | 513096 | 2990 | 0.014489939 |
| LINE/L1 | 281597 | 553 | 0.007952359 |
| Low_complexity | 277765 | 116 | 0.007844142 |
| Satellite | 250944 | 551 | 0.007086712 |
| DNA/TcMar-Tigger | 221606 | 307 | 0.0062582 |
| LTR/Gypsy | 218317 | 455 | 0.006165318 |
| SINE/tRNA | 193505 | 429 | 0.005464622 |
| SINE/5S | 179476 | 1296 | 0.00506844 |
| DNA/hAT-Tip100 | 133264 | 197 | 0.003763404 |
| LINE/RTE-BovB | 128208 | 401 | 0.003620621 |
| PLE/Chlamys | 83874 | 200 | 0.002368619 |
| LINE/I-Jockey | 82930 | 291 | 0.002341961 |
| SINE/tRNA-RTE | 81408 | 54 | 0.002298979 |
| SINE? | 69626 | 101 | 0.001966253 |
| LINE/CR1-Zenon | 49947 | 499 | 0.001410514 |
| SINE/ID | 39799 | 582 | 0.001123932 |
| LINE/L1-Tx1 | 27852 | 197 | 0.000786546 |
| SINE/U | 16878 | 427 | 0.000476638 |
| SINE/5S-Deu-L2 | 5082 | 137 | 0.000143517 |
| snRNA | 2742 | 38 | 7.74347E-05 |
| DNA/Maverick | 2440 | 39 | 6.89061E-05 |
| LINE/L2 | 2029 | 53 | 5.72994E-05 |
| LINE | 2015 | 36 | 5.6904E-05 |
| DNA/hAT-Charlie | 1824 | 42 | 5.15101E-05 |
| LINE/I | 679 | 9 | 1.91751E-05 |
| LTR/Copia | 368 | 7 | 1.03924E-05 |
| DNA/CMC-EnSpm | 268 | 7 | 7.56838E-06 |

| | | | |
|---|---|---|---|
| DNA/PIF-Harbinger | 244 | 6 | 6.89061E-06 |
| LTR/ERV | 166 | 3 | 4.68788E-06 |
| LINE/R2 | 94 | 2 | 2.65458E-06 |
| DNA/hAT | 30 | 1 | 8.47206E-07 |
| total | 341901263 | 476769 | 9.655363889 |

### Section 3: Genomic analyses results

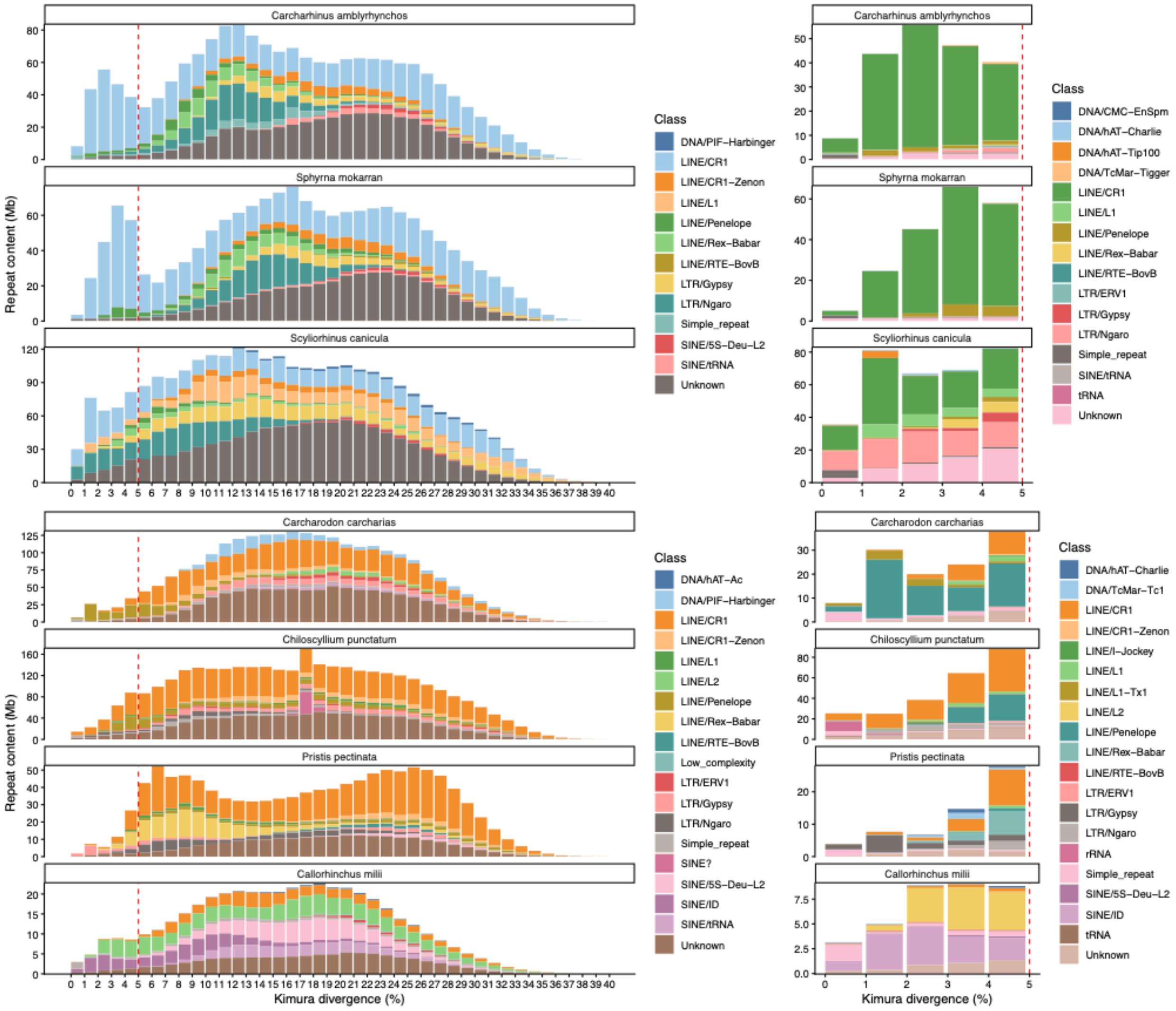

**Figure S4.** Kimura divergence of repeat content by class across seven cartilaginous fish species in order of divergence from *Galeocerdo cuvier*. Right: Full Kimura divergence spectrum across whole genome. Left: Low divergence repeat content (<5%).

### Section 4: Methylation analysis

**Table S5.** Sequencing statistics for methylation calling from PacBio HiFi and Oxford Nanopore Technologies Ultra-long (UL) data generated for the sGalCuv2 reference assembly. All ONT data called using the HAC model.

| Data | Mean depth | Called sites | | | Sites >5X | | Sites >10X | |
|---|---|---|---|---|---|---|---|---|
| | | N sites | Mean depth | Av. meth % | N sites (% retained) | Av. meth % | N sites (% retained) | Av. meth % |
| **HiFi** | 59.54 | 58,939,899 | 61.67 | 63.01 | 58,833,037 (99.8%) | 62.99 | 58,113,229 (98.5%) | 62.88 |
| **ONT 5mCG** | 8.84 | 58,336,541 | 5.46 | 61.60 | 34,590,918 (59.3%) | 63.42 | 4,298,212 (7.4%) | 58.23 |
| **ONT 5hmCG** | | | | 4.21 | | 4.07 | | 3.45 |
| **ONT 5mC** | 8.82 | 1,531,305,930 | 3.12 | 4.14 | 307,752,197 (20.1%) | 2.60 | 6,897,679 (0.5%) | 1.40 |
| **ONT 5hmC** | | | | 0.16 | | 0.08 | | 0.06 |

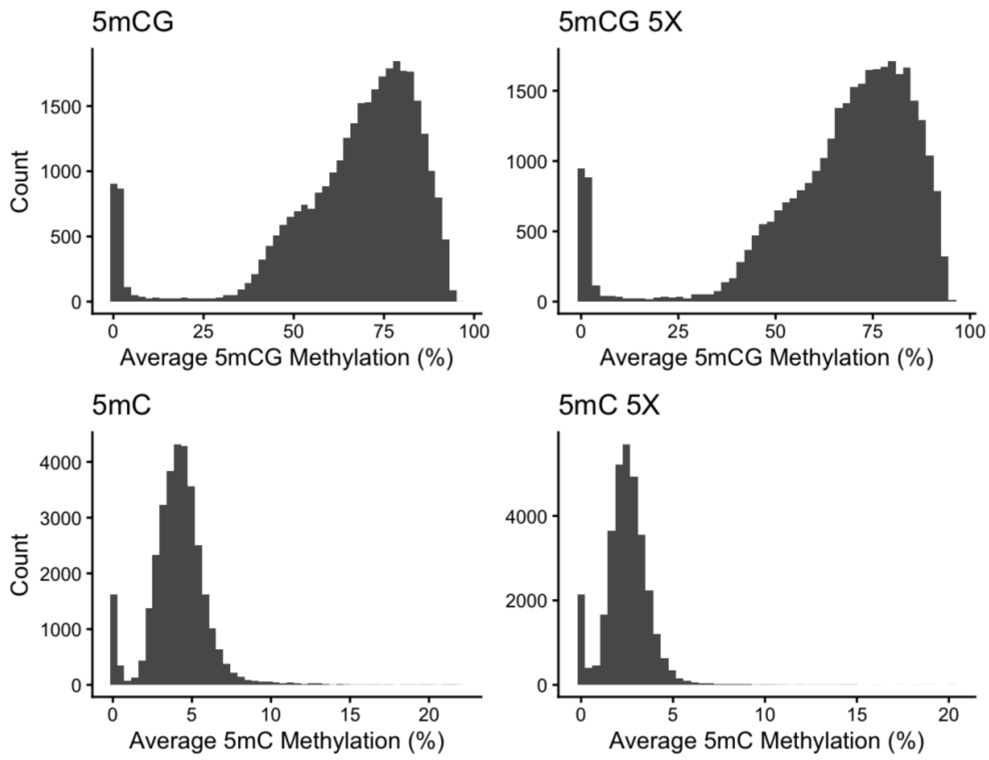

**Figure S5.** Methylation distribution shown as the average percentage of methylation sites across 100KB windows calculated from ONT data. All methylation was estimated using the HAC model, with either the 5mCG tag (top: CpG sites only) or 5mC tag (bottom: all cytosines). Distribution after filtering for sites with at least 5X coverage also shown (right side).

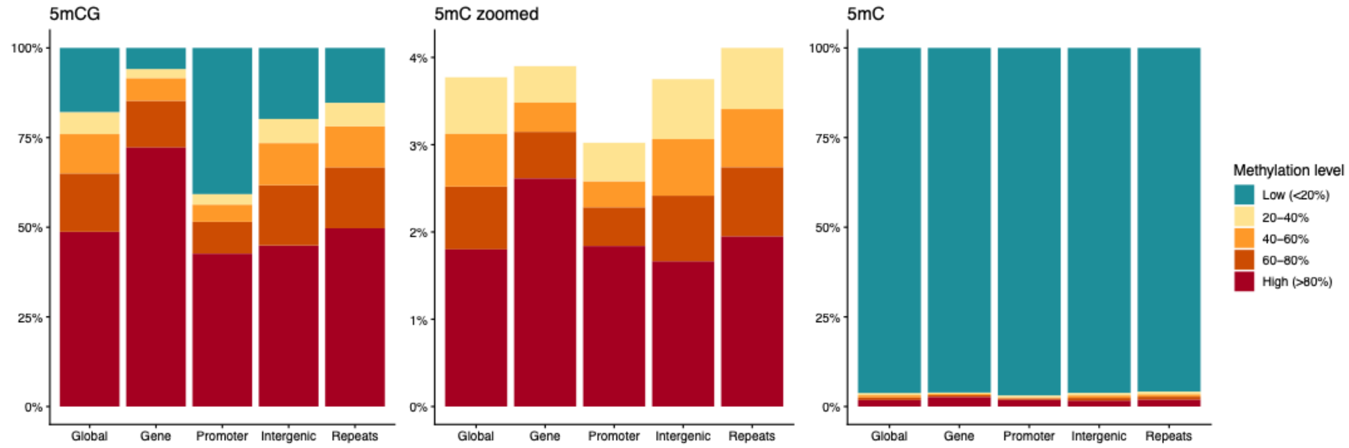

**Figure S6.** Average methylation for genomic regions calculated from ONT data. All methylation was estimated using the HAC model and filtered for sites with at least 5X coverage, with either the 5mCG tag (left: CpG sites only) or 5mC tag (right: all cytosines).

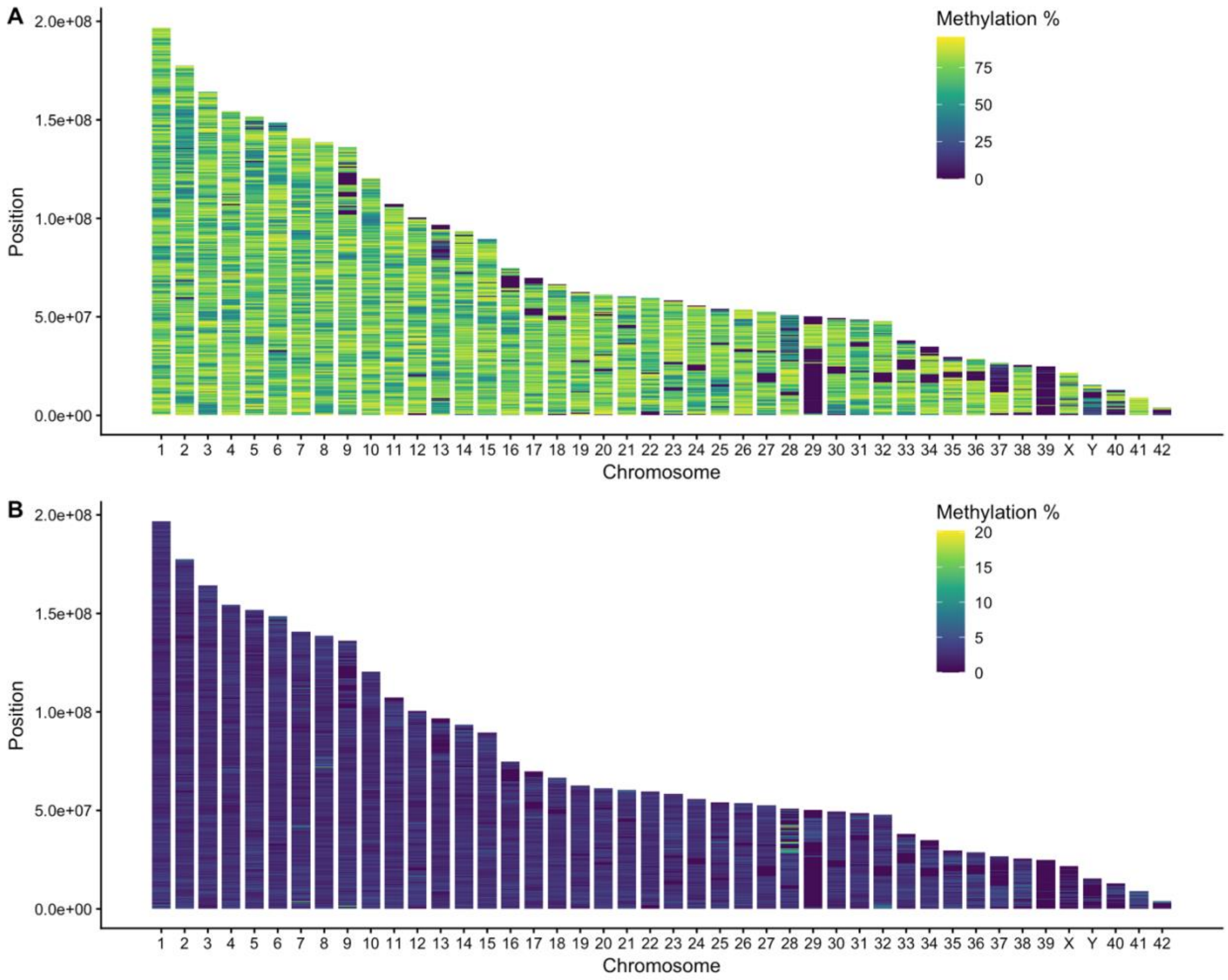

**Figure S7.** Genome-wide methylation across chromosomes, calculated from ONT HAC model for 100KB windows across all chromosomes ordered by length and filtered for a minimum of 5X coverage per site. A) 5mCG B) 5mC

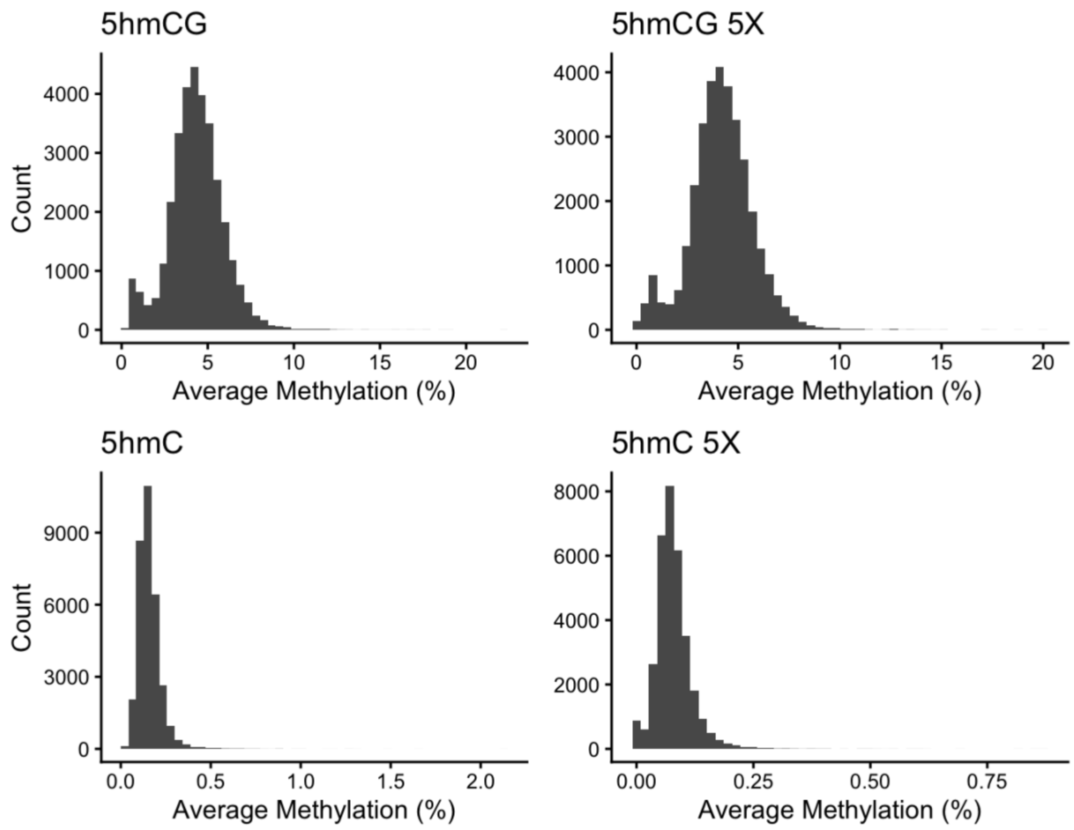

**Figure S8.** Hydroxymethylation distribution shown as the average percentage of methylation sites across 100KB windows calculated from ONT data. All methylation was estimated using the HAC model, with either the 5hmCG tag (top: CpG sites only) or 5hmC tag (bottom: all cytosines). Distribution after filtering for sites with at least 5X coverage also shown (right side).

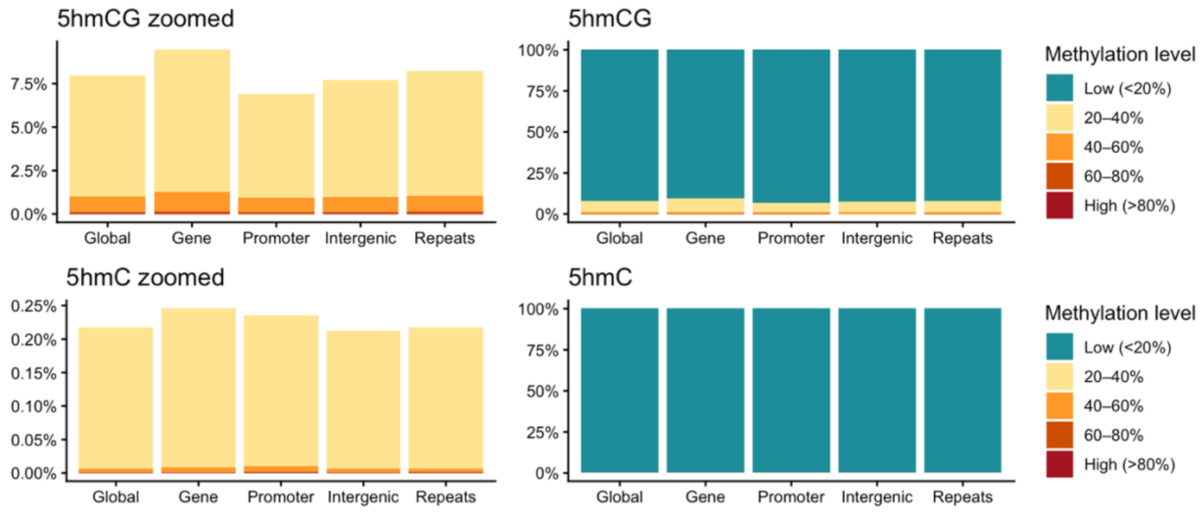

**Figure S9.** Average hydroxymethylation for genomic regions, calculated from ONT data. All methylation was estimated using the HAC model and filtered for sites with at least 5X coverage, with either the 5hmCG tag (left: CpG sites only) or 5hmC tag (right: all cytosines).

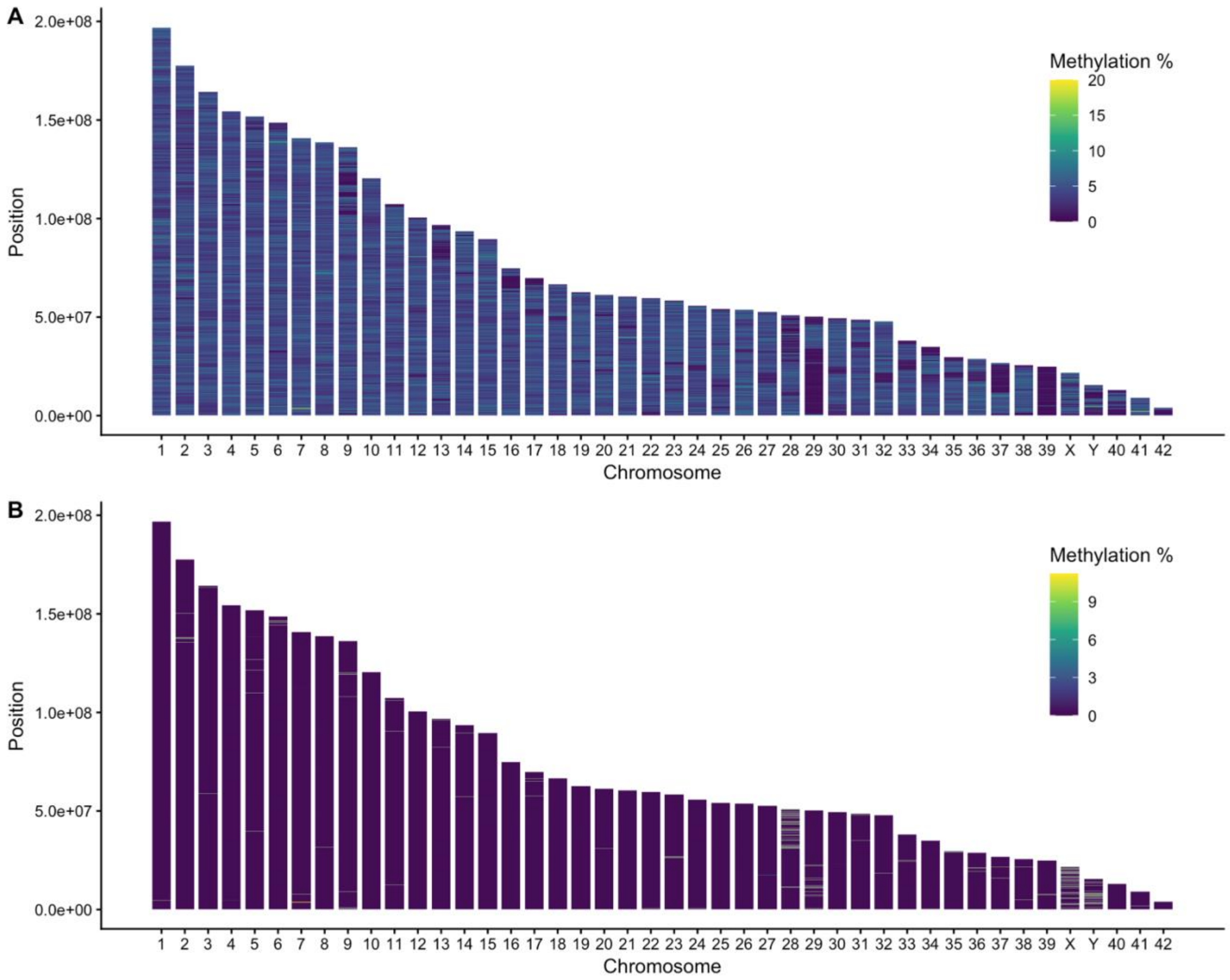

**Figure S10.** Average hydroxymethylation methylation across chromosomes, calculated from ONT HAC model for 100KB windows across all chromosomes ordered by length and filtered for a minimum of 5X coverage per site. A) 5hmCG B) 5hmC
